# Single cell RNA sequencing of calvarial and long bone endocortical cells

**DOI:** 10.1101/849224

**Authors:** Ugur M. Ayturk, Joseph P. Scollan, Alexander Vesprey, Christina M. Jacobsen, Paola Divieti Pajevic, Matthew L. Warman

## Abstract

Single cell RNA-seq (scRNA-seq) is emerging as a powerful technology to examine transcriptomes of individual cells. We determined whether scRNA-seq could be used to detect the effect of environmental and pharmacologic perturbations on osteoblasts. We began with a commonly used *in vitro* system in which freshly isolated neonatal mouse calvarial cells are expanded and induced to produce a mineralized matrix. We used scRNA-seq to compare the relative cell type abundances and the transcriptomes of freshly isolated cells to those that had been cultured for 12 days *in vitro*. We observed that the percentage of macrophage-like cells increased from 6% in freshly isolated calvarial cells to 34% in cultured cells. We also found that *Bglap* transcripts were abundant in freshly isolated osteoblasts but nearly undetectable in the cultured calvarial cells. Thus, scRNA-seq revealed significant differences between heterogeneity of cells *in vivo* and *in vitro*. We next performed scRNA-seq on freshly recovered long bone endocortical cells from mice that received either vehicle or Sclerostin-neutralizing antibody for 1 week. Bone anabolism-associated transcripts were also not significantly increased in immature and mature osteoblasts recovered from Sclerostin-neutralizing antibody treated mice; this is likely a consequence of being underpowered to detect modest changes in gene expression, since only 7% of the sequenced endocortical cells were osteoblasts, and a limited portion of their transcriptomes were sampled. We conclude that scRNA-seq can detect changes in cell abundance, identity, and gene expression in skeletally derived cells. In order to detect modest changes in osteoblast gene expression at the single cell level in the appendicular skeleton, larger numbers of osteoblasts from endocortical bone are required.

## INTRODUCTION

New technologies enable investigators to characterize the transcriptomes of thousands of individual cells in a single experiment[1-3]. These technologies are collectively referred to as single cell RNA sequencing (scRNA-seq) and have been used to determine the diversity of cell types in a complex tissue[1, 4-7], identify novel cell types [8, 9], and identify cells at different stages of differentiation[10-13]. We determined the utility of using scRNA-seq to detect changes in the abundance, differentiation, and transcriptional activity of cells recovered from neonatal mouse calvarium and adult mouse long bone endosteum.

The neonatal mouse calvarium is enriched for bone-forming osteoblasts and their progenitors. A traditional approach in studying osteoblast biology has therefore been to harvest and culture calvarial cells. When supplemented with pro-osteogenic media containing ascorbic acid and β-glycerophosphate, these cells follow a well-defined differentiation cascade towards a mature osteoblast phenotype *in vitro*[14-16]. We performed scRNA-seq on freshly isolated calvarial cells that had been cultured and induced to mineralize *in vitro* to identify similarities and differences between calvarial cells *in vivo* and *in vitro*.

Sclerostin (encoded by the gene *SOST*) is highly expressed by osteocytes and inhibits WNT-signaling by binding to the receptors LRP5 and LRP6 [17, 18]. In humans, *SOST* deficiency causes two high bone mass disorders Sclerosteosis and Van Buchem disease [19, 20], which are recapitulated by *Sost*-null mice [21]. The administration of an antibody that neutralizes Sclerostin induces new bone formation in multiple species, including humans[22-26]. Consequently, an anti-Sclerostin monoclonal antibody has recently been approved by United States Food and Drug Administration to treat patients with osteoporosis[27]. By performing bulk RNA sequencing on mRNA recovered from mouse cortical long bone, we observed that mice with deficient WNT-signaling (i.e., *Lrp5*^-/-^) had a lower abundance of transcripts associated with bone anabolism (e.g., *Col1a1, Col1a2*, and *Bglap*) compared to controls. Conversely, we observed increased bone anabolic transcripts in bulk RNA recovered from the cortical long bones of mice that received Sclerostin-neutralizing antibody (SclAbIII) [28, 29]. Because bulk RNA sequencing cannot determine if a difference in transcript abundance is caused by a change in the number of osteoblasts or a change in the activity of individual osteoblasts, we investigated whether scRNA-seq could detect a change in the number of osteoblasts and/or a change in an osteoblasts’ activity.

## METHODS

### Animals

All animal experiments were approved by the Institutional Animal Care and Use Committees at Boston Children’s Hospital, Massachusetts General Hospital and Weill Cornell Medical College. The experimental design is depicted in Figure 1. For calvarial cell isolation (Figure 1a), twelve P4 pups born to a CD-1 dam (CRL stock #2200) were euthanized by hypothermia and decapitation. For long bone endocortical cell isolation, 11-week-old male C57Bl/6J mice were purchased from The Jackson Labs (Stock # 000664) and acclimatized for 1 week under standard housing conditions. Four mice were then given 2 subcutaneous injections, 3 days apart, of PBS and 4 mice were similarly given subcutaneous injections of SclAbIII (Amgen, Thousand Oaks, CA; 25 mg/kg in PBS). Four days after the 2^nd^ injection the animals were euthanized by CO_2_ inhalation. (Figure 1b).

**Figure 1:**
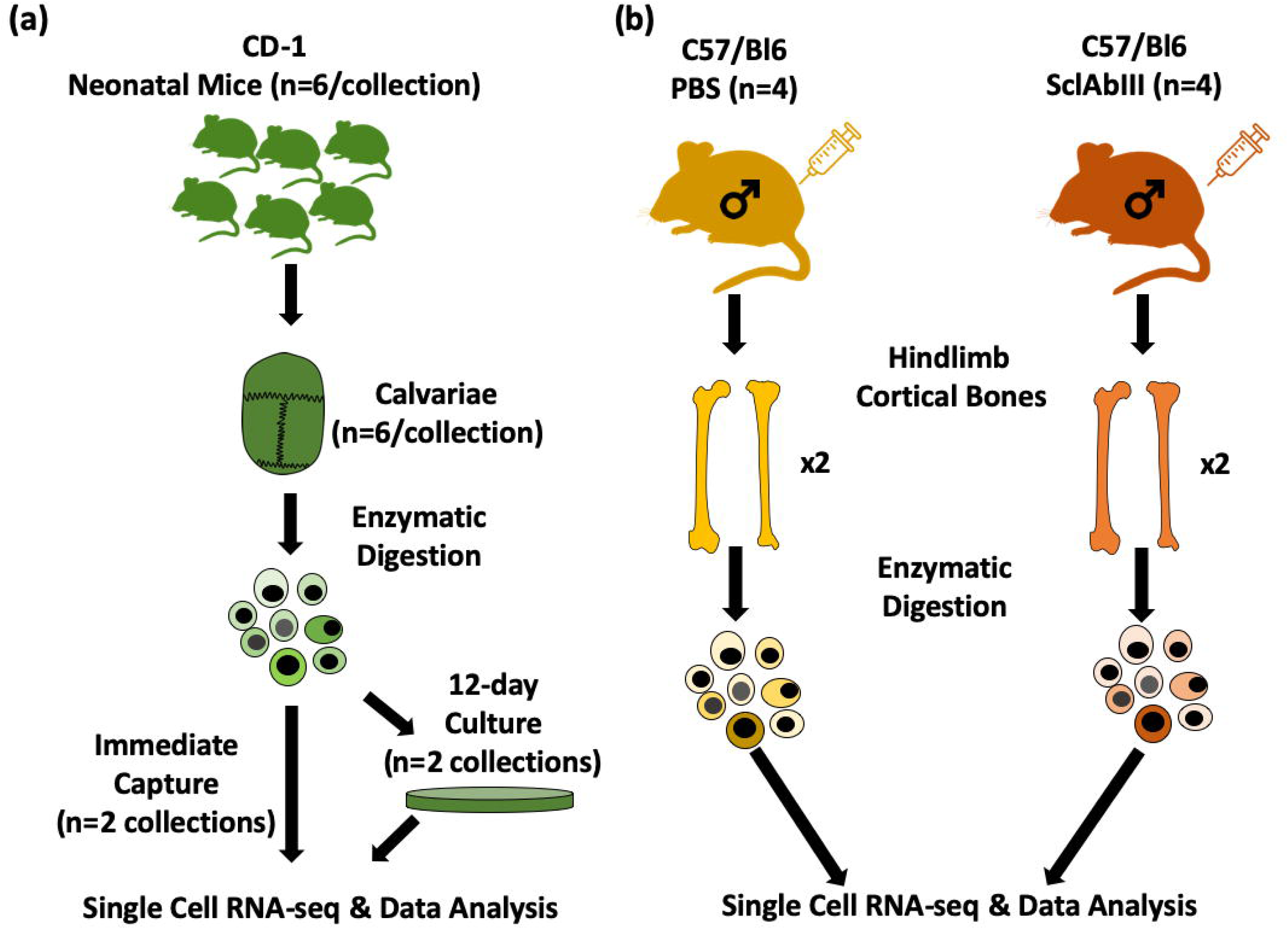
Single cell RNA-seq study design. **(a)** Two sets of samples, each composed of cells that had been isolated CD-1 neonatal mouse calvarium by collagenase-digestion and EDTA treatment (6 calvaria/sample), were collected. An aliquot from each sample was immediately subjected to single cell RNA-seq; the remaining cells were plated, allowed to reach confluence, and cultured under osteogenic conditions for 12 days prior to undergoing single cell RNA-seq. **(b)** 12-week-old C57Bl/6J male mice received 2 SclAbIII (n=4) or 2 PBS (n=4) injections, 3 days apart. Four days later, tibial and femoral diaphyses were collected from each mouse and the endocortical cells were recovered by collagenase digestion.

### Calvarial Cell Isolation

Calvaria were recovered from P4 pups immediately after decapitation. Two samples, each consisting of calvaria from 6 pups were digested in 10 ml collagenase solution (1 mg/ml Collagenase type I and type II (1:3 ratio, Worthington Biochemical Corp., Lakewood, NJ), 1 mM CaCl_2_, 0.1% BSA, 25mM HEPES in αMEM) with gentle agitation at 37°C inside a 5% CO_2_ incubator. The collagenase was replaced every 15 minutes for 5 cycles, followed by 5 mM EDTA (with 0.1% BSA in PBS) treatment for 1 cycle, collagenase for 1 cycle and 5 mM EDTA for the final cycle. Cells released into the medium during the final collagenase cycle and EDTA cycle were combined, mixed with an equal volume of cell culture medium (αMEM + 10% FBS + 1% anti-mycotic) and centrifuged at 500g for 10 min at 4°C. Pelleted cells were re-suspended in cell culture medium and an aliquot from each sample was used to create an scRNA-seq library. The remaining cells in each sample were seeded in 6-well plates at 50,000 cells/well and allowed to reach confluence, while changing the media every 1-2 days. Cells reached confluence at day 5, and were then washed with PBS, separated from the plate via trypsin treatment, transferred to collagen-coated plates (Corning, Corning, NY), and supplemented with 50 μg/ml ascorbic acid and 1 mM β-glycerophosphate for another 7 days in order to initiate osteogenic differentiation. After a total of 12 days in culture, cells were recovered by collagenase digestion for 30 min at 37°C, pelleted and resuspended in cell culture medium, and used to create scRNA-seq libraries. Samples comprised originally of calvaria from 6 pups each were used to create n=2 freshly isolated calvarial cell and n=2 cultured calvarial cell scRNA-seq libraries.

### Endosteal Cell Isolation

Femora and tibiae were collected immediately after euthanasia. The distal and proximal epiphyses were cut away with a scalpel and bones were centrifuged at 13,000 rpm for 1 minute at room temperature to remove bone marrow (Supplementary Figure 1). Periosteum was aseptically removed with a scalpel and the bone was bisected lengthwise to expose the endosteum. Tibiae and femora from each mouse were combined and incubated with 4 ml of collagenase solution (3 mg/ml Collagenase Type IV in PBS, Worthington Biochemical Corp., Lakewood, NJ) inside a 5% CO_2_ chamber at 37°C for 15 min under continuous agitation. Cells recovered from this initial digestion were discarded since they contained a large fraction of red blood cells (Supplementary Figure 2). The long bones were digested in two additional 4 ml collagenase solutions for 30 minutes each, and the cells recovered with each digestion were mixed with an equal volume of α-MEM (w/10%FBS + 1% anti-mycotic) and combined. After pelleting the cells at 2,000 x g for 5 minutes, the pellet was resuspended in 0.7 ml of PBS (w/ 0.04% BSA) and then used to prepare scRNA-seq libraries. To verify that collagenase digestion efficiently removed cells from the endosteal bone surface, the bones were fixed in 10% formalin, decalcified with 10% EDTA, processed and embedded in paraffin, sectioned and stained with hematoxylin and eosin (H&E), and compared to similar sections from bones that had been placed in PBS (Supplementary Figure 3).

### Single Cell Capture and RNA-seq Library Preparation

We used the Chromium Single Cell 3’ system (10X Genomics, Pleasanton, CA) to capture and sequence single cell mRNA, following the manufacturer’s instructions. We loaded ∼10,000 cells per endosteal specimen and ∼6,000 cells per calvarial specimen. Individual cells were captured inside oil droplets along with barcoding beads, such that the mRNA contents of each cell were ligated with cell-specific oligo-barcodes (Supplementary Figure 1). Following single cell capture, barcoded mRNA and reverse-transcription solutions were immediately transferred to a thermal cycler, wherein barcoded cDNA was generated. The pooled cDNAs were then chemically fragmented, ligated with sequencing adapters, amplified and purified with magnetic beads (Beckman Coulter, Brea, CA). Quality checks for libraries were performed with gel electrophoresis and TapeStation analysis (Agilent Technologies, Santa Clara, CA).

### Sequencing and Data Analysis

Single cell RNA-seq libraries were pooled and run on the Illumina NextSeq platform. Data analysis was performed using the Cellranger and Seurat pipelines [30]. Briefly, raw sequence data were de-multiplexed into specimen-specific bins, and mapped to the mouse genome (mm10) with STAR aligner[31]. The mapped sequence data and the associated unique molecular identifiers (UMI) were used to determine the number of captured cells, and the transcriptome of each cell.

We excluded cells that had higher-than-expected mitochondrial transcripts and transcriptional diversity (indicated by the number of unique transcripts per cell). We combined the fresh calvarial cell and cultured calvarial cell datasets, performed tSNE analysis [1, 30], and used the unbiased cluster-detection algorithm of Seurat[30] to identify transcriptionally distinct cell populations. We performed similar analyses with the SclAbIII- and PBS-treated endocortical cell datasets. We regressed cell-cycle associated transcriptional signals from our data, as cells going through mitotic division might be registered as distinct populations. We then identified the transcripts that set each cell population apart from the others. We quantified the number of cells in each cell population, in a sample-specific manner. We also quantified the mRNA expression levels in each cell population with respect to their sample of origin, and ultimately the treatment group of origin.

### Differential Gene Expression Analysis Between Long Bone Endocortical Cell Specimens

We performed differential expression analysis between the SclAbIII- and PBS-treated groups, in a cell cluster-specific manner, using edgeR[32]. We determined significance with p < 0.05 after correction for multiple hypothesis testing. As scRNA-seq can detect a limited portion of each cell’s transcriptome, we imposed an additional detectability threshold, such that > 50% of the cells in either group had to have > 0 expression of the tested gene.

## RESULTS

### Calvarial Cell Recovery and Sequencing

We collected ∼3 million cells from each sample of 6 pooled calvaria, aliquots of which were used to generate scRNA-seq libraries. We obtained an average of ∼194 million reads from the fresh and cultured calvarial single cell libraries, which represented ∼75,000 reads/cell, and ∼2,522 cells per library following pre-processing and filtering.

Seurat’s unbiased cluster detection algorithm detected 11 distinct cell populations within the freshly collected calvarial cells and 8 distinct cell populations within the cultured calvarial cells (Figure 2a). The majority of the fresh calvarial cell population consisted of mesenchymal cells (clusters #3, #4, #7, #10 and #17, which represented >75% of all cells in each sample). Clusters #3, #4 and #7 exhibited transcriptional gradients in osteoblast markers (e.g., *Col1a1, Bglap* and *Dmp1*), suggesting they represent osteoblasts at different stages of differentiation. Clusters #10 and #17 exhibited transcriptional profiles compatible with chondrocytes and alpha smooth muscle actin-expressing (αSMA+) smooth muscle cells, respectively (Figure 2b). The remaining fresh calvarial cell clusters expressed transcripts found in endothelial cells (*Pecam1/Cd31*, #18), red-blood-cells (*Hbb-bs*, #12 and #15), granulocytes (#14), B-cells (#16) and myeloid cells (#13).

**Figure 2:**
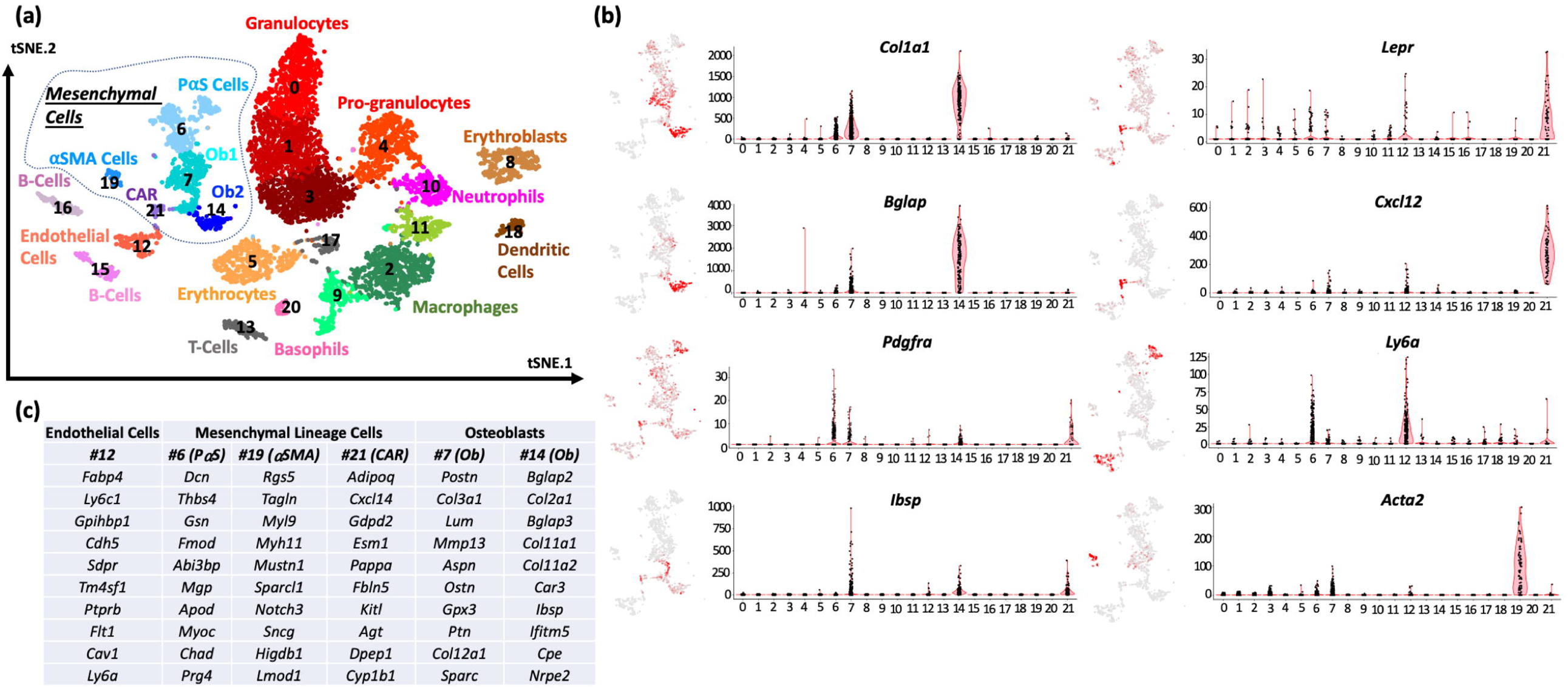
Cell clusters identified among cultured and fresh calvarial cells. **(a)** tSNE plot wherein each dot represents a single cell, and closely grouped cells indicate transcriptionally similar cells. Cultured and freshly isolated cells form separate clusters (dashed line). Of note, cells in cluster #8 express transcripts associated with mesenchymal (e.g. *Col1a1, Acta2*) and hematopoietic (*Ptprc* and *Csf1r*) lineages, have higher than expected UMI counts, and likely represent artifact due to 2 cell types being captured in a single droplet. **(b)** Heatmaps of gene expression (left) and associated violin plots (right) indicate cluster-specific expression of representative genes. Osteoblast-lineage cells among freshly isolated calvarial cells are distinctly marked by the expression of *Mfap4* (#3), *Postn* (#4) and *Bglap* (#7). Similarly, cultured osteoblast-lineage cells express *Acta2* and *Col1a1* (#1, 2, 5 and 9), whereas hemaotopoietic cells express *Csf1r* and *Ptprc* (#0, 6 and 8). **(c)** Top 10 transcript markers corresponding to each cell cluster as determined by Seurat (OB: osteoblast).

### scRNA-seq Indicates that Calvarial Cell Populations and Transcriptomes Change in vitro

Cultured calvarial cells clustered separately from freshly isolated cells (Figure 2a). Within the cultured calvarial cell populations, clusters #1, #2, #5 and #9 grouped near one-another and expressed transcripts (e.g. *Col1a1, Runx2, Sp7* and *Alpl*) that suggest they are mesenchymal cells differentiating along the osteoblast lineage. However, in contrast to the freshly isolated calvarial cells, no cultured calvarial cell cluster expressed transcripts typically associated with mature osteoblasts, such as *Bglap, Dmp1* or *Ifitm5*. Also, whereas only 10 and 12% of freshly isolated calvarial cell samples appeared hematopoietic in origin (clusters #12 to #16, as indicated by the expression of transcripts including *Csf1r* and *Ptprc*), this percentage increased to 45 and 48% in the cultured cells (clusters #0 and #6), respectively. These data suggest that hematopoietic cells proliferate more or survive better than mesenchymal cells in culture.

### Long Bone Endocortical Cell Recovery and Sequencing

H & E staining indicated that our collagenase/EDTA cell recovery method removed nearly all endosteal surface cells, but did not remove embedded osteoblasts or osteocytes (Supplementary Figure 3). We recovered ∼1 million cells from the pooled tibiae and femora of individual mice. We generated ∼ 65 million reads per scRNA-seq library, which after deconvolution, alignment, and cell-specific gene expression represented an average of ∼47,000 reads/cell, and 1053 cells/mouse following filtering for outlier cells based on mitochondrial and unique transcript content.

Seurat’s unbiased cluster detection algorithm defined 22 cell populations within the long bone endocortical samples of PBS- and SclAbIII-treated mice (Figure 3a). We utilized the single cell transcriptome database (specifically bone marrow scRNA-seq data) developed by the Tabula Muris Consortium [33] to assign identities to these populations. Three cell clusters (#6, #7, and #14) from the long bone endocortical cell libraries, representing ∼13% of the cells, had transcriptional profiles consistent with their being mesenchymal cells and osteoblasts (Figure 3b). The most abundant cell population among these, cluster #6 contains peri-arteriole stromal cells (also known as PαS cells[34]), which express *Col1a1, Pdgfra* and *Ly6a/Sca1*. Clusters #7 and #14, representing ∼7% of the sequenced endocortical cells in each animal, are contiguous and express osteoblast-associated transcripts (e.g., *Bglap, Ifitm5*, and *Dmp1*). Differences in expression between these two clusters suggest cluster #14 comprises more mature osteoblasts (Figure 3b).

**Figure 3:**
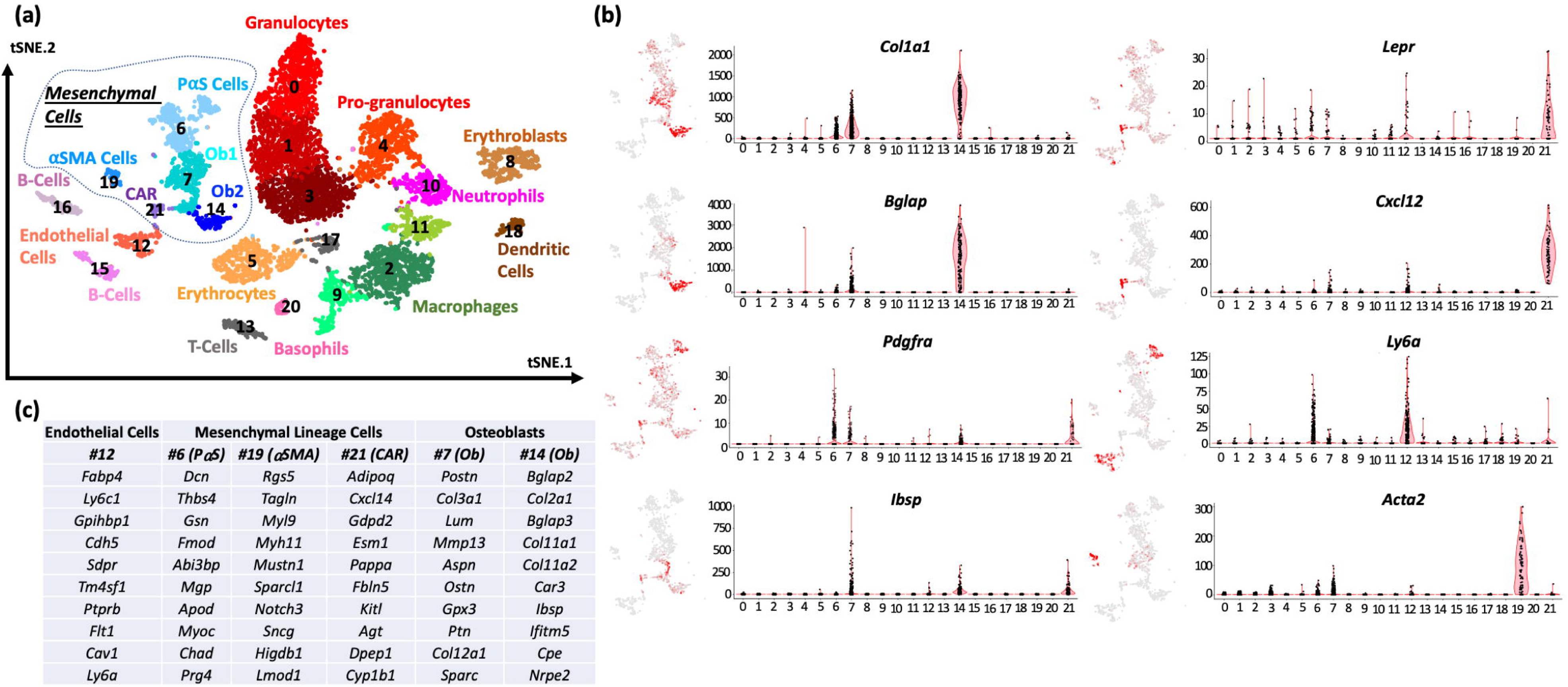
Endosteal cell clusters detected by scRNA-seq. **(a)** tSNE plot wherein each dot represents a single cell, and closely grouped cells indicate transcriptionally similar cells. Five clusters of cells (highlighted inside the dashed line) represent mesenchymal progenitors and osteoblasts at different stages of differentiation, as indicated by their individual expression profiles depicted in (b) and (c). **(b)** Heatmaps of gene expression (left) and associated violin plots (right) indicate cluster-specific expression of representative genes. Increasing expression of *Col1a1* and *Bglap* across clusters #7 and #14 suggest that these clusters represent osteoblasts. Cells in cluster #6 are marked by *Pdgfra* and *Ly6a*/*Sca1* expression, cluster #21 is marked by *Lepr* and high levels of *Cxcl12* expression, cluster #12 is marked by *Ly6a/Sca1* expression, and cluster #19 is marked by *Acta2/aSMA* expression. **(c)** Top 10 transcript markers corresponding to each of the mesenchymal and endothelial cell clusters as determined by Seurat (Ob: Osteoblast)

In marked contrast to freshly recovered calvarial cells, the majority of cells (>80%) that we recovered from cortical long bone were hematopoietic in origin. Four clusters, representing ∼ 40% of the cells, express transcripts associated with granulocytes and pro-granulocytes. Six clusters, representing ∼30% of the cells express transcripts associated with T-cells, B-cells, neutrophils, basophils, dendritic cells and macrophages. Two clusters, representing ∼12% of the cells, express transcripts associated with erythroblasts and erythrocytes.

Two other previously described cell types, Cxcl12-Abundant-Reticular (CAR) cells [35] and peri-arteriole smooth muscle (αSMA+) cells [36] comprise clusters #21 and #19, respectively (Figure 3c). Interestingly, cells in cluster #21 express transcripts seen in mature osteoblasts (e.g., *Ibsp, Serpinh1/Hsp47, Gja1/Connexin43, Pcolce*), but they do not express *Col1a1* or *Bglap* at a level detectable by scRNA-seq (Supplementary Table 1).

### No Discernable Effect of SclAbIII Treatment on Endocortical Cells Measured by scRNA-seq

There was no discernable difference in the relative proportions of the different cell clusters from the long bones of PBS and SclAbIII-treated mice (Figure 4a). We did not identify mesenchymal cells undergoing cell-cycling in either group. We also did not observe a significant difference in the single cell expression levels of osteoblast-associated transcripts such as *Col1a1, Bglap*, and *Cpz* in clusters #7 or #14 between PBS- and SclAbIII-treated mice (Figure 4b). Instead, we only detected significantly altered expression of 9 protein-coding, non-ribosomal, transcripts in cluster #7 (*Hint1, Oaz1, Fos, H3f3b, Cst3, Tceb2, Sep15, Dynll1* and *Omd*) and 5 protein-coding, non-ribosomal, transcripts in cluster #14 (*Cst3, Fau, Ftl1, Gpx3* and *Loxl2*) following SclAbIII treatment.

**Figure 4:**
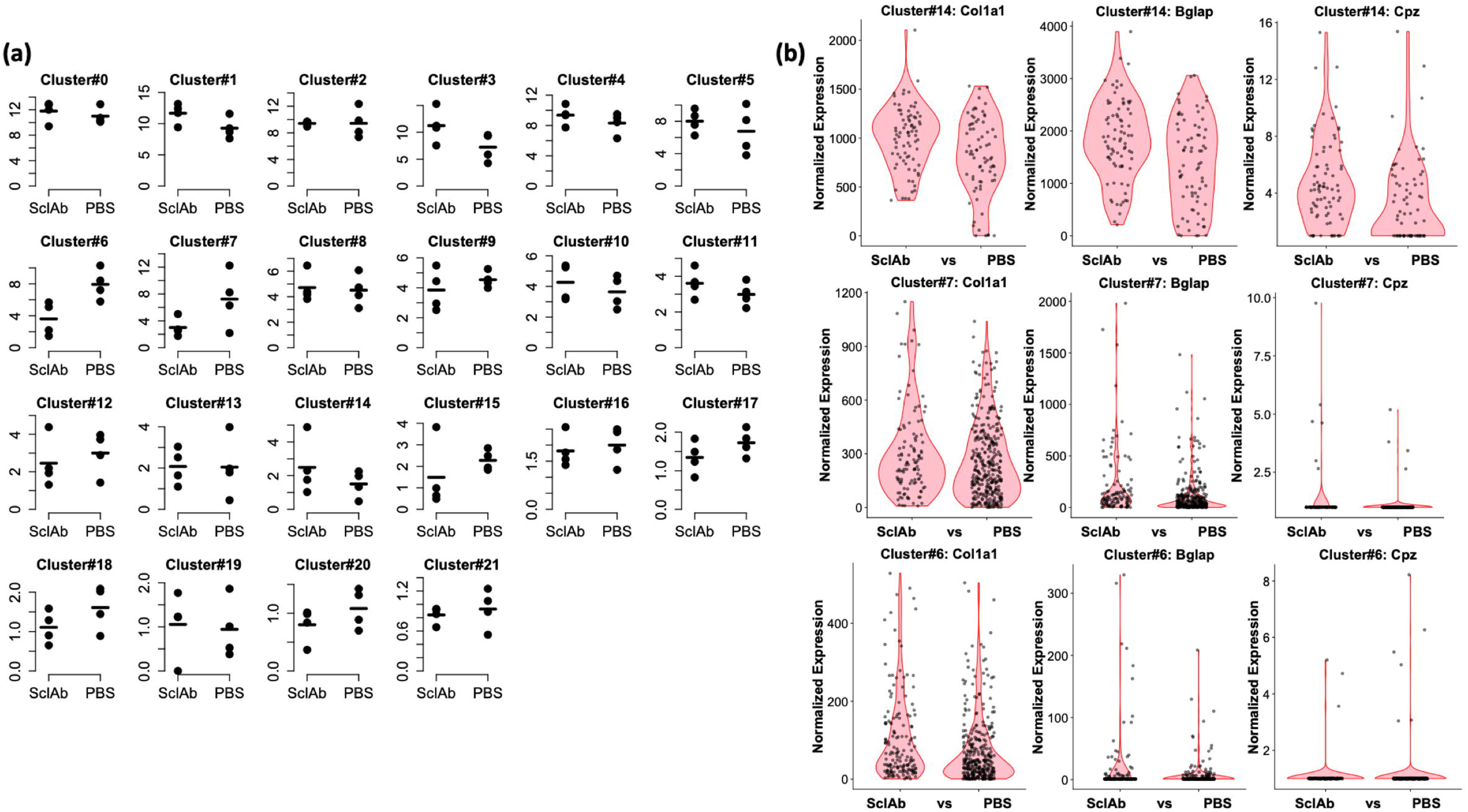
Reproducibility of endocortical scRNA-seq data across biologic replicates. **(a)** Dot plots depict the percentage of cells in each cluster across endocortical cells collected from 8 mice (n=4 SclAbIII-treated and n=4 PBS-treated), suggesting high reproducibility in the quantification of transcriptionally distinct cells. **(b)** Violin plots show no significant difference in the expression of 3 genes associated with osteoblast anabolism (*Col1a1, Bglap*, and *Cpz*) due to SclAbIII treatment in the mesenchymal or osteoblast cell clusters.

### Osteoblasts Freshly Isolated from Calvaria and Cortical Long Bone Have Similar Transcriptomes

We observed that freshly isolated calvarial osteoblasts had different transcriptomes from those of cultured calvarial osteoblasts (Figure 2a). This led us to determine whether there may also be differences between freshly isolated osteoblasts from calvaria and cortical long bone. Each scRNA-seq library contained clusters of cells that expressed low or high levels of *Bglap*, which we used to define early and late stage osteoblasts, respectively. When we compared the transcriptomes of osteoblasts from calvaria and long bone, we found a strong concordance (Figure 5).

**Figure 5:**
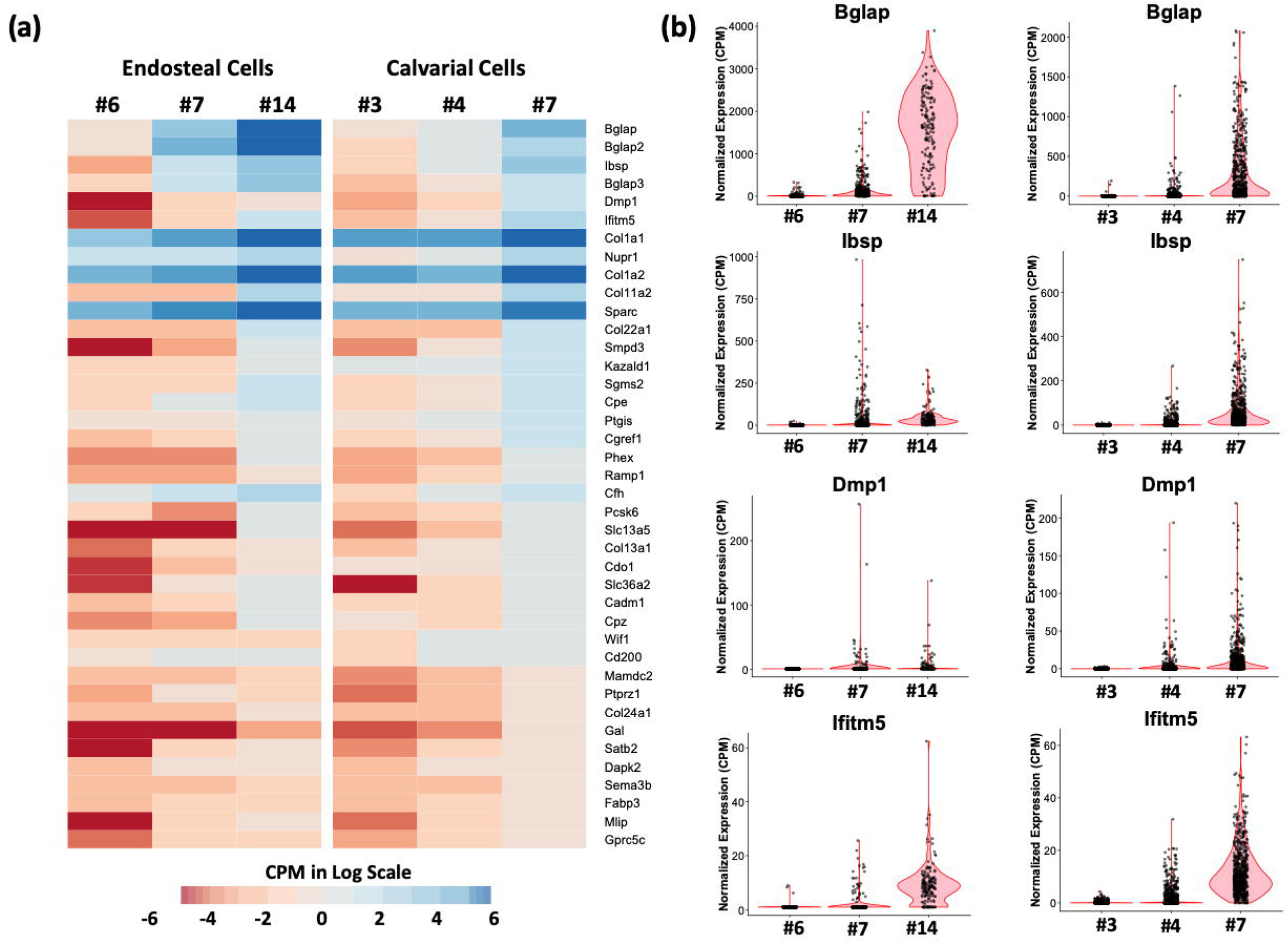
High concordance between the transcriptomes of freshly isolated calvarial and long bone endocortical osteoblasts. **(a)** Heatmap depicts the mean expression levels of the top 40 gene markers for calvarial osteoblast cluster #7, across the osteoblast-lineage cells in endosteal and calvarial collections. **(b)** Violin plots depict the changes in the expression of top osteoblast markers *Bglap, Ibsp, Dmp1* and *Ifitm5* across the long bone endosteal (left) and calvarial (right) cell clusters.

### Robustness of Bone-derived Single Cell RNA-seq Data

Obtaining endocortical cells from 8 individual mice allowed us to assess the consistency with which we recovered different cell types and to determine whether gene expression within a cell type correlated between animals. We observed that the relative proportions of cell types were similar across all animals and there was a high Pearson correlation coefficient between the transcriptomes of each cluster across animals (Supplementary Figure 4). The only exceptions to these observations are for clusters #19 and #21 where lower correlations across samples were observed (R^2^ range: 0.37-0.79), likely due to the low number of cells (<90 cells) that were contained in these clusters. Although we only had 2 samples to compare for the freshly obtained calvarial cells, these samples also were closely matched, and we calculated high Pearson correlation coefficients when transcriptomes of each cluster were compared between samples. (Supplementary Figure 5).

Using whole tissue RNA-seq, we had previously identified a set of transcripts, whose abundances were dramatically elevated when long bone tissue was exposed to collagenase[28]. These transcripts included *Fos, Fosb, Jun, Il6*, possibly indicating a stress response. When we interrogated our scRNA-seq data, we found that these stress response transcripts were at low or undetectable levels (Supplementary Figures 6 and 7, Supplementary Table 1). We infer that the cells expressing these stress-response transcripts were not sampled by scRNA-seq, most likely because cells that remained embedded express these transcripts.

## DISCUSSION

Single cell RNA-seq can identify discrete cell types in cultured cells and in complex tissues, based on each individual cell’s transcriptome. The ability to interrogate the transcriptomes of large numbers of individual cells has led to the discovery of previously unrecognized cell types [8, 9] and has detected changes in cellular diversity in response to genetic or environmental perturbation[37-39]. Recovering individual cells from skeletal tissue is more challenging than from cultured cells or from other tissues whose cells can be easily separated by brief digestion and/or physical disruption. For this reason, *in vitro* studies of osteoblast differentiation and mineralization typically expand cells in culture and then chemically induce matrix mineralization[15, 16]. In order to study freshly isolated osteoblasts or their precursors, skeletal tissues need to be subjected to sequential enzymatic digestions for an hour or longer [14, 15, 40]. We employed scRNA-seq to assess similarities and differences between osteoblasts that were freshly obtained from calvaria versus long bone endocortices, between fresh calvarial cells and those that had been expanded in culture, and between long bone endocortical cells from PBS- and SclAbIII-treated animals. We observed greater enrichment for osteoblasts that had been freshly isolated from calvaria compared to those that had been freshly isolated from long bone endocortices. More than 40% of calvarial cells had transcriptomes expected of osteoblasts, but only 7% of endocortical cells exhibited these profiles (Figures 2a and 3a). Reassuringly, however, the transcriptomes of freshly isolated calvarial and endocortical osteoblasts were highly similar, suggesting that osteoblast precursors converge to a shared transcriptional phenotype during differentiation, regardless of their origin (Figure 5). Although we cannot exclude the possibility that many of these similarities are solely due to collagenase induced transcription, we think this is unlikely since most of the transcripts used to define these cells as osteoblasts are present in cortical bone bulk RNA-seq data and are seen in osteoblasts *in situ* [28]. Finally, we observed high Pearson correlation coefficients within individual cell populations from 2 freshly collected or cultured calvarial specimens and 8 endocortical specimens, indicating that we are reproducibly recovering cells (Supplementary Figures 4-5 and 8-9).

Calvarial cells collected after 120 minutes (i.e. 8 cycles) of enzymatic digestion from P4 mice are enriched for chondrocytes, osteoblasts and their precursors. In our hands, ∼6% of the freshly isolated cells are macrophages (cluster #13, Figure 2a), based on the expression of genes such as *Ptprc/Cd45, Cd68* and *Csf1r*. However, after we expanded the calvarial cells in culture and induced differentiation, we found that 34% of the cells were macrophages (clusters #0 and #6). Our results are consistent with those of Chang et al., who describe a population of cells they termed “osteomacs,” which accounted for ∼16% of freshly collected calvarial cells and proliferated in culture [41]. Colony stimulating factor 1 (CSF1) had previously been proposed as a mediator of signaling between osteoblasts and osteomacs [42, 43]; consistent with this model, *Csf1* and *Csf1r* expression distinctly mark cells we transcriptionally identified as osteoblasts and macrophages, respectively. Moreover, our scRNA-seq data identifies other cell type-specific transcripts (such as *Cd68, Cd14, Ccl9*) that may also be involved in paracrine signaling. We did not detect *Bglap*-expressing cells among the calvarial osteoblasts that had been expanded in culture and induced to differentiate. This contrasts with freshly isolated calvarial cells that exhibited abundant *Bglap* expression. These data are consistent with previous studies that indicate freshly isolated calvarial osteoblasts initially de-differentiate *in vitro* [14, 44].

Although we previously used bulk RNA-seq of cortical bone to observe significant increases in transcripts associated with bone anabolism in mice given a short course of SclAbIII [29], we did not observe an increase using scRNA-seq in our long bone endocortical samples. Several factors may account for this discordance. First, we were underpowered to observe modest increases in gene expression using scRNA-seq. Bulk RNA-seq sampled millions of cells per mouse and detected the expression of ∼10,000 genes. In contrast, scRNA-seq sampled only ∼75 osteoblasts/mouse from n=4 mice/group and detected the expression of ∼1,500 transcripts on average (Supplementary Figure 10). Therefore, we likely sampled too few mice to be able to detect meaningful changes in cell type abundance, and too few cells to be able to detect meaningful changes in gene expression. Second, we administered 4 doses of neutralizing antibody for our bulk RNA-seq studies but only 2 doses for the scRNA-seq study. Finally, our scRNA-seq data was derived primarily from isolated osteoblasts, whereas the bulk RNA-seq data was derived from specimens containing osteocytes.

In conclusion, we have employed scRNA-seq to show that freshly recovered osteoblasts from newborn calvaria and adult endocortical long bone have highly correlated transcriptomes, that freshly isolated calvarial cells undergo changes in their relative abundances and transcriptomes when expanded and differentiated *in vitro*, and that scRNA-seq is not yet as sensitive for detecting changes in osteoblast gene expression as bulk RNA-seq. As better methods for recovering and enriching for osteoblasts from endocortical samples are developed, and greater depths of coverage for transcripts from individual cells are obtained, we anticipate scRNA-seq will become a useful tool for monitoring the effects of genetic, environmental, and pharmacologic perturbations on endocortical bone cells.

## ACKNOWLEDGMENTS

This study was funded by grants P30 AR066261, R01 AR053237, R01 AR064231 and R21AR067388 from NIAMS.

## SUPPLEMENTARY FIGURES

**Supplementary Figure 1: Single cell RNA-seq workflow for endocortical and calvarial bone specimens.**

**Supplementary Figure 2: Two-fold reduction in bulk *Hbb* expression between endosteal cell collections #1 and #2, measured by qRT-PCR.**

**Supplementary Figure 3: Reduction of endocortical surface cells following collagenase treatment.** We found that there were very few cells left on the endosteal surface of bone samples treated with collagenase, whereas there were numerous clusters of cells on PBS treated surfaces (white arrows).

**Supplementary Figure 4: Reproducibility of cluster-specific gene expression measurements in specimens from SclAbIII- (top) and PBS-treated mice (bottom).** We calculated the mean gene expression profile in a cell cluster- and mouse-specific manner, and calculated R^2^ values by performing intra-group comparisons (top: SclABIII, bottom: PBS) between each experimental mouse. Our cell cluster-specific measurements were highly reproducible among the majority of the clusters, except for #19 and #21. The intra-group variability of gene expression in these clusters is likely due to low number of cells (<90 cells total in each cluster).

**Supplementary Figure 5: Reproducibility of cluster-specific gene expression measurements in biologic replicates of fresh (highlighted with blue) and cultured (highlighted with red) cells.**

**Supplementary Figure 6: Violin plots depicting the cluster-specific expression profile of the top 8 genes previously found to be induced by collagenase treatment (Ayturk et al., 2013) in endosteal specimens.** Note that these genes are expressed at either undetectable (as in the case of *Il6* and *Ftl2*) or low to moderate levels in all cells (as reference, mean *Bglap* and *Col1a1* expression levels were found to be 876 and 490 in Cluster#14, respectively).

**Supplementary Figure 7: Violin plots depicting the cluster-specific expression profile of the top 8 genes previously found to be induced by collagenase treatment (Ayturk et al., 2013) in calvarial cells.** Note that these genes are expressed at either undetectable (as in the case of *Ftl2*) or low to moderate levels in all cells.

**Supplementary Figure 8: Reproducibility of scRNA-seq on calvarial cells indicated by tSNE distribution.**

**Supplementary Figure 9: Reproducibility of scRNA-seq on endocortical cells indicated by tSNE distribution.**

**Supplementary Figure 10: The number of UMIs and genes detected per cell in each endosteal cell cluster.** We found that the the magnitude of transcriptional diversity was highly variable across the 22 cell populations we identified. *Hbb*+ red blood cells (cluster#8) were the least diverse, whereas mature osteoblasts (cluster#14), neutrophils (cluster#10) and macrophages (cluster#11) expressed up to ∼3,000 distinct genes per cell.

## SUPPLEMENTARY TABLES

**Supplementary Table 1: Mean gene expression levels per cell cluster in calvarial cells.**

**Supplementary Table 2: Mean gene expression levels per cell cluster in endocortical cells.**

## REFERENCES

1. Macosko, E.Z., et al., Highly Parallel Genome-wide Expression Profiling of Individual Cells Using Nanoliter Droplets. Cell, 2015. 161(5): p. 1202–1214.

2. Zheng, G.X., et al., Massively parallel digital transcriptional profiling of single cells. Nat Commun, 2017. 8: p. 14049.

3. Zilionis, R., et al., Single-cell barcoding and sequencing using droplet microfluidics. Nat Protoc, 2017. 12(1): p. 44–73.

4. Baryawno, N., et al., A Cellular Taxonomy of the Bone Marrow Stroma in Homeostasis and Leukemia. Cell, 2019. 177(7): p. 1915–1932 e16.

5. Shekhar, K., et al., Comprehensive Classification of Retinal Bipolar Neurons by Single-Cell Transcriptomics. Cell, 2016. 166(5): p. 1308–1323 e30.

6. Tikhonova, A.N., et al., The bone marrow microenvironment at single-cell resolution. Nature, 2019. 569(7755): p. 222–228.

7. Wolock, S.L., et al., Mapping Distinct Bone Marrow Niche Populations and Their Differentiation Paths. Cell Rep, 2019. 28(2): p. 302–311 e5.

8. Debnath, S., et al., Discovery of a periosteal stem cell mediating intramembranous bone formation. Nature, 2018. 562(7725): p. 133–139.

9. Plasschaert, L.W., et al., A single-cell atlas of the airway epithelium reveals the CFTR-rich pulmonary ionocyte. Nature, 2018. 560(7718): p. 377–381.

10. Chan, M.M., et al., Molecular recording of mammalian embryogenesis. Nature, 2019. 570(7759): p. 77–82.

11. Kanton, S., et al., Organoid single-cell genomic atlas uncovers human-specific features of brain development. Nature, 2019. 574(7778): p. 418–422.

12. Pijuan-Sala, B., et al., A single-cell molecular map of mouse gastrulation and early organogenesis. Nature, 2019. 566(7745): p. 490–495.

13. Popescu, D.M., et al., Decoding human fetal liver haematopoiesis. Nature, 2019.

14. Zhong, Z.A., N.J. Ethen, and B.O. Williams, Use of Primary Calvarial Osteoblasts to Evaluate the Function of Wnt Signaling in Osteogenesis. Methods Mol Biol, 2016. 1481: p. 119–25.

15. Lian, J.B. and G.S. Stein, Development of the osteoblast phenotype: molecular mechanisms mediating osteoblast growth and differentiation. Iowa Orthop J, 1995. 15: p. 118–40.

16. Lynch, M.P., et al., The influence of type I collagen on the development and maintenance of the osteoblast phenotype in primary and passaged rat calvarial osteoblasts: modification of expression of genes supporting cell growth, adhesion, and extracellular matrix mineralization. Exp Cell Res, 1995. 216(1): p. 35–45.

17. Li, X., et al., Sclerostin binds to LRP5/6 and antagonizes canonical Wnt signaling. J Biol Chem, 2005. 280(20): p. 19883–7.

18. Semenov, M., K. Tamai, and X. He, SOST is a ligand for LRP5/LRP6 and a Wnt signaling inhibitor. J Biol Chem, 2005. 280(29): p. 26770–5.

19. Balemans, W., et al., Increased bone density in sclerosteosis is due to the deficiency of a novel secreted protein (SOST). Hum Mol Genet, 2001. 10(5): p. 537–43.

20. Brunkow, M.E., et al., Bone dysplasia sclerosteosis results from loss of the SOST gene product, a novel cystine knot-containing protein. Am J Hum Genet, 2001. 68(3): p. 577–89.

21. Li, X., et al., Targeted deletion of the sclerostin gene in mice results in increased bone formation and bone strength. J Bone Miner Res, 2008. 23(6): p. 860–9.

22. Li, X., et al., Sclerostin antibody treatment increases bone formation, bone mass, and bone strength in a rat model of postmenopausal osteoporosis. J Bone Miner Res, 2009. 24(4): p. 578–88.

23. Li, X., et al., Inhibition of sclerostin by monoclonal antibody increases bone formation, bone mass, and bone strength in aged male rats. J Bone Miner Res, 2010. 25(12): p. 2647–56.

24. Marenzana, M., et al., Sclerostin antibody treatment enhances bone strength but does not prevent growth retardation in young mice treated with dexamethasone. Arthritis Rheum, 2011. 63(8): p. 2385–95.

25. Ominsky, M.S., et al., Two doses of sclerostin antibody in cynomolgus monkeys increases bone formation, bone mineral density, and bone strength. J Bone Miner Res, 2010. 25(5): p. 948–59.

26. Padhi, D., et al., Single-dose, placebo-controlled, randomized study of AMG 785, a sclerostin monoclonal antibody. J Bone Miner Res, 2011. 26(1): p. 19–26.

27. Markham, A., Romosozumab: First Global Approval. Drugs, 2019. 79(4): p. 471–476.

28. Ayturk, U.M., et al., An RNA-seq protocol to identify mRNA expression changes in mouse diaphyseal bone: applications in mice with bone property altering Lrp5 mutations. J Bone Miner Res, 2013. 28(10): p. 2081–93.

29. Kedlaya, R., et al., Sclerostin inhibition reverses skeletal fragility in an Lrp5-deficient mouse model of OPPG syndrome. Sci Transl Med, 2013. 5(211): p. 211ra158.

30. Butler, A., et al., Integrating single-cell transcriptomic data across different conditions, technologies, and species. Nat Biotechnol, 2018. 36(5): p. 411–420.

31. Dobin, A., et al., STAR: ultrafast universal RNA-seq aligner. Bioinformatics, 2013. 29(1): p. 15–21.

32. Robinson, M.D., D.J. McCarthy, and G.K. Smyth, edgeR: a Bioconductor package for differential expression analysis of digital gene expression data. Bioinformatics, 2010. 26(1): p. 139–40.

33. Tabula Muris, C., et al., Single-cell transcriptomics of 20 mouse organs creates a Tabula Muris. Nature, 2018. 562(7727): p. 367–372.

34. Morikawa, S., et al., Prospective identification, isolation, and systemic transplantation of multipotent mesenchymal stem cells in murine bone marrow. J Exp Med, 2009. 206(11): p. 2483–96.

35. Zhou, B.O., et al., Leptin-receptor-expressing mesenchymal stromal cells represent the main source of bone formed by adult bone marrow. Cell Stem Cell, 2014. 15(2): p. 154–68.

36. Matthews, B.G., et al., Osteogenic potential of alpha smooth muscle actin expressing muscle resident progenitor cells. Bone, 2016. 84: p. 69–77.

37. Klaus, J., et al., Altered neuronal migratory trajectories in human cerebral organoids derived from individuals with neuronal heterotopia. Nat Med, 2019. 25(4): p. 561–568.

38. Sharir, A., et al., A large pool of actively cycling progenitors orchestrates self-renewal and injury repair of an ectodermal appendage. Nat Cell Biol, 2019. 21(9): p. 1102–1112.

39. Szczerba, B.M., et al., Neutrophils escort circulating tumour cells to enable cell cycle progression. Nature, 2019. 566(7745): p. 553–557.

40. Greenblatt, M.B., et al., The Unmixing Problem: A Guide to Applying Single-Cell RNA Sequencing to Bone. J Bone Miner Res, 2019. 34(7): p. 1207–1219.

41. Chang, M.K., et al., Osteal tissue macrophages are intercalated throughout human and mouse bone lining tissues and regulate osteoblast function in vitro and in vivo. J Immunol, 2008. 181(2): p. 1232–44.

42. Alexander, K.A., et al., Osteal macrophages promote in vivo intramembranous bone healing in a mouse tibial injury model. J Bone Miner Res, 2011. 26(7): p. 1517–32.

43. Winkler, I.G., et al., Bone marrow macrophages maintain hematopoietic stem cell (HSC) niches and their depletion mobilizes HSCs. Blood, 2010. 116(23): p. 4815–28.

44. Jonason, J.H. and R.J. O’Keefe, Isolation and culture of neonatal mouse calvarial osteoblasts. Methods Mol Biol, 2014. 1130: p. 295–305.

